# Quantitative Cellular-Resolution Map of the Oxytocin Receptor in Postnatally Developing Mouse Brains

**DOI:** 10.1101/719229

**Authors:** Kyra T. Newmaster, Zachary T. Nolan, Uree Chon, Daniel J. Vanselow, Abigael R. Weit, Manal Tabbaa, Shizu Hidema, Katsuhiko Nishimori, Elizabeth A.D. Hammock, Yongsoo Kim

## Abstract

Oxytocin receptor (OTR) plays critical roles in social behavior development. Despite its significance, brain-wide quantitative understanding of OTR expression remains limited in postnatally developing brains. Here, we validated and utilized fluorescent reporter mice (OTR^venus/+^) to examine OTR cells across postnatal periods. We developed postnatal 3D template brains to register whole brain images with cellular resolution to systematically quantify OTR cell densities. We found that cortical regions showed temporally and spatially heterogeneous patterns with transient postnatal OTR expression without cell death. Cortical OTR cells were largely not GABAergic neurons with the exception of cells in layer 6b. Subcortical regions showed similar temporal regulation except the hypothalamus. Moreover, our unbiased approach identified two hypothalamic nuclei with sexually dimorphic OTR expression. Lastly, we created a website to easily share our imaging data. In summary, we provide comprehensive quantitative data to understand postnatal OTR expression in the mouse brain.

## Introduction

Oxytocin receptor (OTR) mediates oxytocin (OT) signaling which plays a critical role in the development of social behavior for animals including humans ^1–3^. Animal models lacking functional OTR show social behavior impairment ^4,5^ suggesting that OTR expression is important for normal social behavior. OTR expression begins early in life with peak cortical OTR expression coinciding with critical postnatal developmental windows for social learning ^2,6,7^ This transient OTR expression in the developing cortex is thought to play an important role in facilitating neural circuit maturation ^8,9^ For instance, OTR in postnatally developing brains has been implicated in multisensory binding ^10^, maturation of GABAergic neurons ^11^, and synapse formation and maturation between neurons ^12,13^.

During the early postnatal period and adulthood, many different brain regions contain OTR expressing cells that are either excitatory or inhibitory neurons ^6,12,14,15^ OTR expressing neurons in different brain regions have been linked to neural circuit specific functions such as facilitating social reward in the ventral tegmental area ^16^, social recognition in the anterior olfactory nucleus^17^, and social memory in the hippocampal CA2 region ^18,19^. However, we have limited knowledge on the temporal and regional expression patterns of OTR throughout the entire brain. Previous studies investigating OTR expression mainly utilized receptor autoradiography binding assays, histological methods (e.g., immunohistochemistry using specific antibodies), or transgenic reporter animals ^6,15,20,2^. Most of these studies, if not all, examined selected brain regions by histological methods, which is difficult to apply for whole brain analysis across developmental periods due to variable staining results, laborious procedures, and semi-quantitative assessment.

To overcome this issue, we developed new postnatal template brains at different postnatal (P) developmental periods (P7, 14, 21, and 28) with detailed anatomical labels based on Allen Common Coordinate Framework ^22^. Then, we expanded our existing quantitative brain mapping platform (qBrain)^23^ to image, detect, and quantify fluorescently labeled cells at the cellular resolution from postnatally developing brains (developmental qBrain; dqBrain). We applied this method to quantify the number and density of OTR (+) cells using OTR-Venus knock-in reporter mice (OTR^venus/+^) after confirming its faithful representation of endogenous OTR expression using fluorescent *in situ* hybridization ^20^. We found temporally and spatially heterogeneous cortical and subcortical expression with early postnatal peak densities. Our cumulative labeling revealed that cortical OTR reduction into adulthood is mainly driven by receptor down-regulation without cell death. Furthermore, we identified sexually dimorphic OTR expression in two hypothalamic nuclei. Lastly, we deposited postnatal template brains and high-resolution image data with user friendly visualization in our website (http://kimlab.io/brain-map/OTR/) to facilitate open data sharing.

## Result

### Choice of fluorescent reporter mice for OTR expression

To quantify OTR expression across the whole brain, we used two transgenic reporter mice that express fluorescent reporters under the OTR promotor. The lines we examined include a BAC transgenic OTR-eGFP reporter mouse ^24^ and a knock-in OTR^venus/+^ heterozygote mouse, called “OTR-Venus” hereafter, that encodes a fluorescent reporter gene (Venus) in place of the genomic OTR coding region^20^. We initially observed significant discrepancies in the number and location of cells reporting OTR expression between the two mouse lines (Figure 1). In order to independently validate these observations, we used single-molecule mRNA fluorescent *in situ* hybridization against OTR in postnatally developing mouse brains. We first confirmed the specificity of the OTR *in situ* hybridization by comparing expression of OTR mRNA wild type littermates mice (WT; n = 3 mice, Figure 1A-J) and OTR knockout mice (OTR^venus/venus^; n = 2 mice, Figure 1K) at P21. OTR^venus/venus^ mice expressed no OTR mRNA whereas their wild-type littermates showed robust expression. Then, we compared our OTR *in situ* hybridization results (WT; n = 3 mice) to fluorescent reporter expression from both OTR-Venus (n = 10) and OTR-eGFP mice (n = 10) at P21. We found that Venus expression from OTR-Venus mice overall matched to endogenous OTR expression very closely while OTR-eGFP often lacked comparable expression (false negative) or misrepresented OTR expression (false positive) in several brain regions (Figure 1A-I). For example, the prelimbic cortex (PL) and the taenia tecta (TT) showed distinct OTR expressions in both OTR *in situ* and OTR-Venus while very little expression in OTR-eGFP mice (Figure 1B, C). We also observed that GFP labeled cells in OTR-eGFP mice were mostly restricted to the superficial layer of the somatosensory cortex while OTR-Venus mice showed a population of Venus-labeled cells in both superficial and deep layer which corresponded to the RNA *in situ* results for analogous areas (Figure 1E). In the posterior cortical area, the OTR-Venus mice exhibit OTR expression that is well-matched to our OTR *in situ* data while OTR-eGFP reports little expression in the layer 2 of the visual cortex (white arrows in Figure 1H). RNA *in situ* data also shows that OTR is strongly expressed in the bed nucleus of stria terminalis posterior interfascicular division (BSTif) which is correctly reported by the OTR-Venus reporter (Figure 1F). However, OTR-eGFP mice report GFP expression in the BST posterior principal nucleus (BSTpr), not in the BSTif (Figure 1F). Moreover, robust OTR expression in the posteromedial cortical amygdala (COApm) was observed in both the OTR *in situ* data and the OTR-Venus reporter while little expression was found in the OTR-eGFP reporter (Figure 1I). We further analyzed OTR-Venus expression at adult stage (at P56) in relation to OTR mRNA expression data from publically available *in situ* database from Allen Institute for Brain Science ^25^ and confirmed comparable expression patterns in OTR-Venus mice (Figure S1). We then examined whether the Venus mRNA and OTR mRNA were co-expressed in the same cells from OTR-Venus mice by using double fluorescent *in situ* hybridization (Figure 1J). We confirmed that the majority of Venus positive cells also express OTR mRNA (83.8%, 321 OTR positive cells among 383 Venus positive cells in the cortex, the amygdala, and the hippocampus, N = 3, P21 mice).

**Figure 1.**
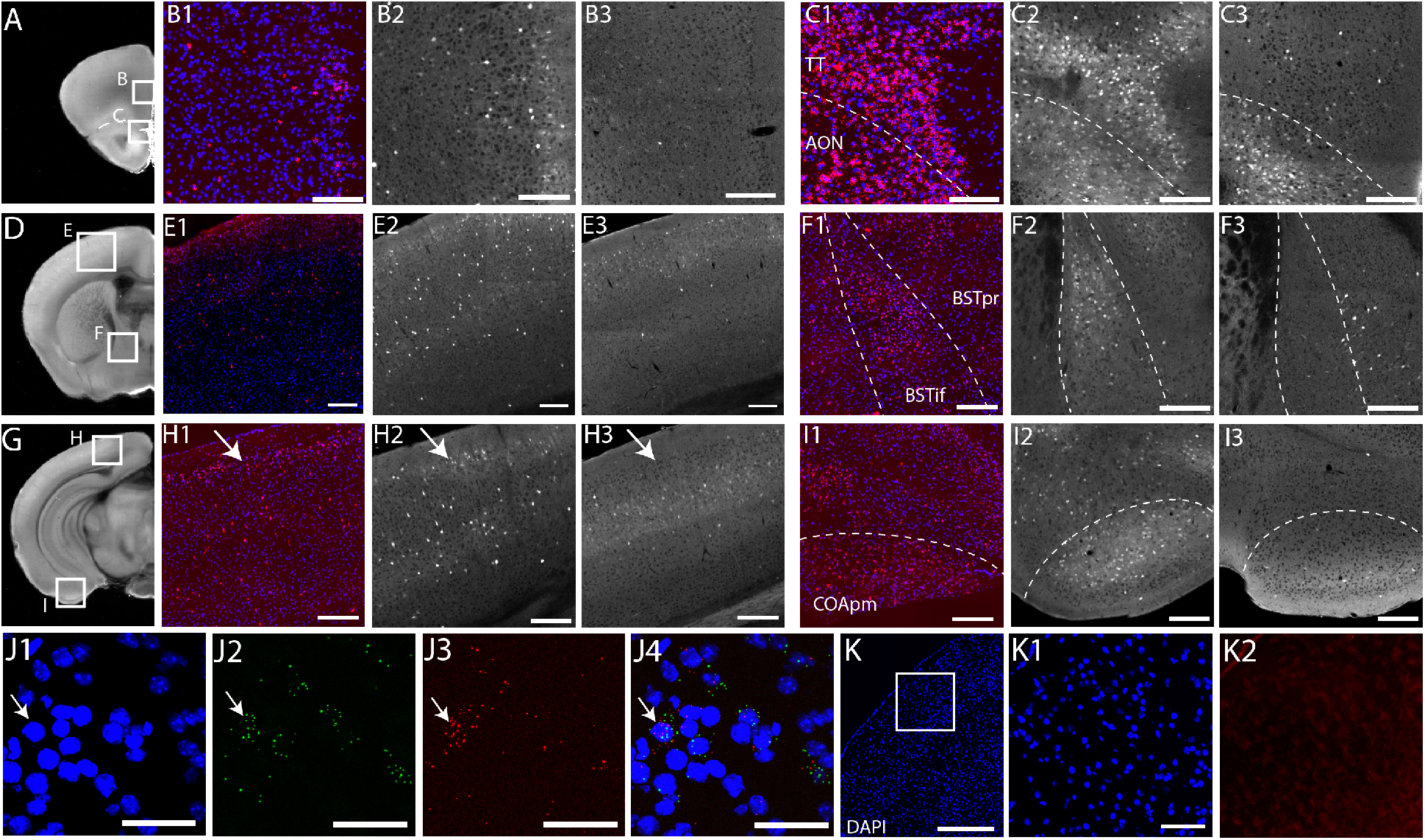
Characterization of OTR transgenic reporter mice. (A-I) Comparison between the OTR fluorescent *in situ* hybridization and OTR transgenic reporter mouse lines at P21. Scale bar = 200 μm. The white boxes in the first column represent brain regions in zoomed-in pictures on subsequent columns. The second and the fifth column for the OTR *in situ*, the third and the sixth for the OTR-Venus mice, and the fourth and the seventh for the OTR-eGFP mice. (B) the prelimbic cortex, (C) the taenia tecta (TT) and the anterior olfactory nucleus (AON), (E) the primary somatosensory cortex, (F) the bed nucleus of stria teminalis (BST) interfascicular (if) and principal (pr) nucleus, (H) the visual cortex, and (I) the cortical amygdala posterior medial (COApm) area. Note the corresponding patterns between the OTR *in situ* and the OTR-Venus, but not OTR-eGFP. (J) Double fluorescent *in situ* against the OTR and the Venus in the cortex from the OTR-Venus mice. (J1) DAPI nuclear staining, (J2) Venus *in situ*, (J3) OTR *in situ*, and (J4) the merged view. The white arrows indicate an example of both OTR and Venus positive cells. Scale bar = 50 μm. (K) OTR *in situ* hybridization on OTR knockout (OTR^venus/venus^) mice. (K1) DAPI staining and (K2) OTR *in situ* in the somatosensory cortex from the white boxed area in K. Note the lack of OTR puncta. Scale bar = 400 μm for (K) and 100 μm for (K1-2).

Collectively, we concluded that the OTR-Venus mice can serve as a good reporter line to examine the developmental trajectory of the OTR expression.

### Quantitative brain-wide mapping pipelines in postnatally developing brains

We previously established a quantitative brain mapping method (termed “qBrain”) that can count the number and the density of fluorescently labeled cells in over 600 different anatomical regions across the entire adult mouse brain with cellular resolution precision ^23^. The method consists of whole brain imaging at cellular resolution using serial two-photon tomography, machine learning based algorithms to detect fluorescently labeled cells, image registration to a reference brain, and statistical analysis ^23^. To extend the method to map signals in postnatally developing brains, we established registration template brains from different early postnatal periods: P7, 14, 21, and 28 (Figure 2) ^26^. First, we chose the best imaged brain at each age (Figure S2). Then, we registered brains from the same age to the initial template brain. Lastly, we averaged all transformed brains to generate an averaged template brain at each age (N = 8 brains at P7, 15 at P14, 12 at P21, and 17 at P28). Furthermore, we generated age-matched anatomical labels by transforming labels from the adult brain based on the Common Coordinate Framework (CCF) from Allen Institute for Brain Science to template brains from younger ages (Figure S2). With these tools, termed “developmental qBrain” (dqBrain), we were able to register our image data to age-matched template brains to quantify fluorescently labeled cells across the entire brain at different postnatal ages (Figure 2, Movie S1-5).

**Figure 2.**
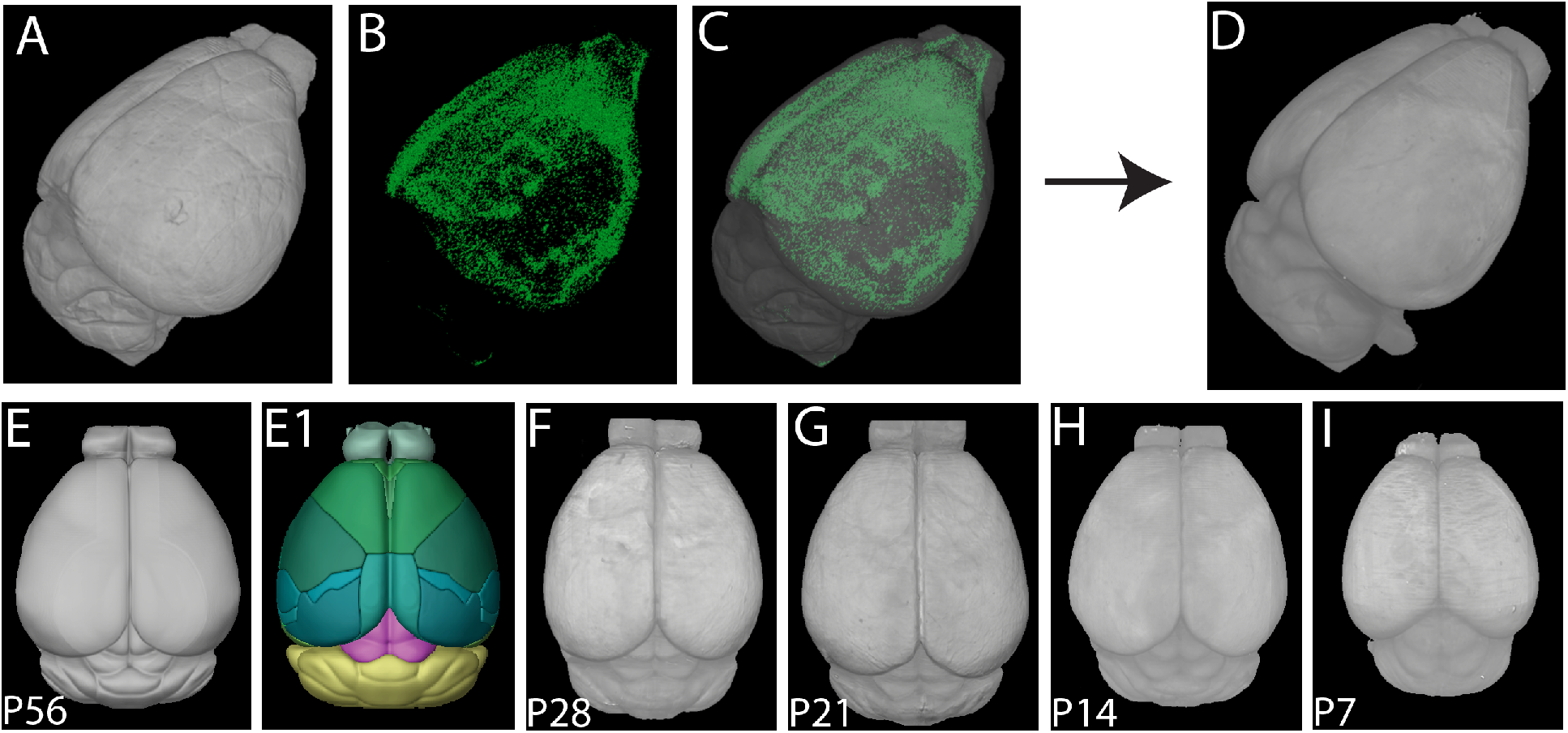
Quantitative brain mapping method to examine OTR expression in developing postnatal mouse brains. (A-C) Reconstructed 3D brains from serial two-photon tomography imaging of the P14 OTR-Venus mouse brain (A), detected Venus positive cells (B), and their overlay (C). (D) The registration template brain for automated cell counting for P14 brains. (E-I) Template brains at different postnatal ages. (E) The adult Allen CCF background template and (E1) its anatomical labels (E1). Newly generated template brains at P28 (F), P21 (G), P14 (H), and P7(I).

### Developmental expression pattern of OTR neurons in the isocortex (neocortex)

To examine regional and temporal heterogeneity of OTR expression, we imaged OTR-Venus mice at five different postnatal days (P7, 14, 21, 28 and 56, N = 5 males and 5 females per age). First, we examined OTR expression in the isocortex (Figure 3, Movie S1-5). Our data showed that overall cortical OTR density reaches its peak at P21 and decreases into adulthood (the red line in Figure 3B). We also noticed spatially heterogeneous expression in different cortical areas (Figure 3B). For example, the anterior cingulate and the retrosplenial cortex, parts of the medial association area, showed low OTR density, while the visual and lateral association areas (e.g., the temporal association area) showed higher OTR density (Figure 3B). Mapping data also revealed temporally heterogeneous patterns. For example, the somatosensory area reached its near peak expression at P14 (bottom panel in Figure 3A; black line in 3B) while the visual area showed rapid increases up to P21 (top panel in Figure 3A; blue line in 3B). To further visualize the temporally and spatially heterogeneous OTR expression patterns more intuitively, we used cortical flatmaps throughout the developing brain^23^. Cortical flatmaps are digitally flattened 2D maps of 3D cortical areas that use evenly spaced bins as a spatial unit to quantify and to display detected signals ^23^. The cortical flatmap clearly highlighted regional differences in OTR developmental expression with early expression in visual, medial prefrontal, and lateral association area as early as P7 (Figure 3C). In contrast, somatosensory regions showed little OTR expression at P7 with a rapid increase in OTR density at P14 (the yellow arrow in the Figure 3C). By P21, regional heterogeneity attained an adult-like pattern although adults showed lower OTR density overall (Figure 3C). Interestingly, the medial prefrontal cortex (mPFC) showed a less dramatic decrease in OTR density as mice progressed to adulthood when compared to other cortical regions, which matches previous results reporting robust OTR expression in the adult mPFC ^14,27^ (the green line in the Figure 3B, the green arrow in 3C).

**Figure 3.**
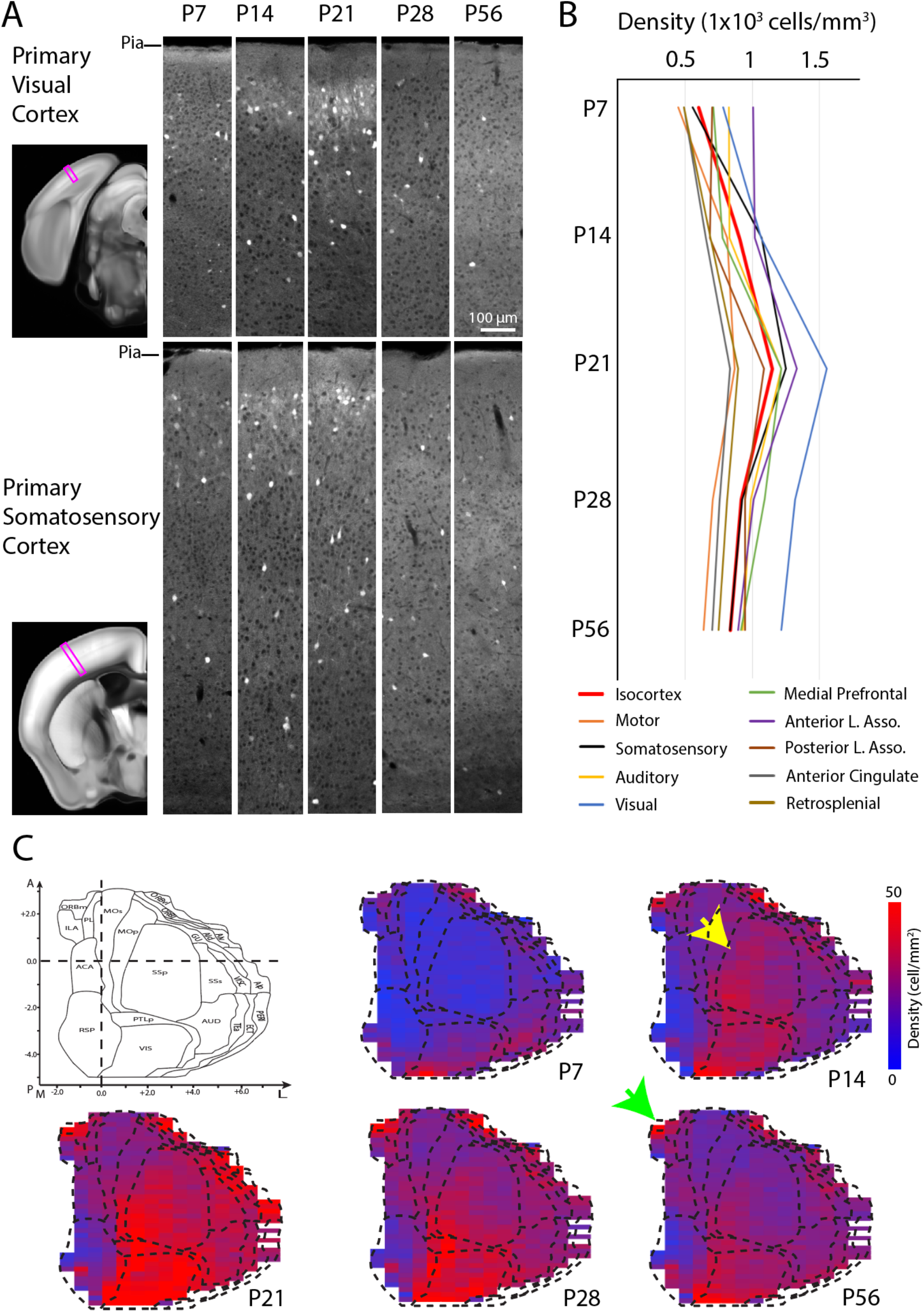
Developmental trajectory of OTR cells in the isocortex from OTR-Venus mice. (A) Representative images from the primary visual and the primary somatosensory cortices (purple boxes) in OTR^Venus/+^ mice at P7, 14, 21, 28, and 56 (columns to the right). Note clustered and dispersed OTR expression in the superficial and deep cortical layers, respectively. (B) Average densities of OTR-Venus cells in different isocortical areas at 5 different postnatal ages. Anterior lateral (L) association (Asso) area for lateral orbital, gustatory, visceral, agranular insular; Posterior L. Asso. for temporal association, perirhinal, ectorhinal cortex. See Table S1 for more details. (C) 2D cortical flatmap representation of OTR-Venus expression pattern at different developmental time points. The heat map represents OTR-Venus densities in evenly spaced bins in the cortical flatmap. Note overall peak expression in all cortical regions at P21. See also Figure S3 for layer specific cortical flatmaps. The yellow at P14 and the green arrow at the P56 flatmap highlights somatosensory cortex and medial prefrontal cortex, respectively. Full name of acronyms can be found in Table S1.

We also noticed higher OTR density in the superficial cortical layers (Layer 1 – 3) compared to deeper layers (Layer 5 – 6) particularly at P14 and P21 (Figure 3A). To understand how this cortical layer specific expression affects developmental OTR patterns, we used layer specific cortical flatmaps to visualize the superficial and deep layer expression patterns separately (Figure S3). We found that OTR in the superficial layer appears earlier and peaks earlier (at around P14) than the deep layers. The superficial layers also show a more pronounced reduction in adulthood compared to the deep layer (Figure S3). At P14, the superficial layer expression patterns of the somatosensory cortex are well-matched with previous OTR autoradiography and mRNA *in situ* data (green arrow in Figure S3A) ^6,10^. Moreover, relative OTR expression in different cortical layers across postnatal periods clearly showed that OTR expression is predominantly found in the superficial layers at P7 and P14 but shifts to similar or even relatively greater density in the deep layer. This pattern is largely driven by the transient OTR expression in the superficial layers (Figure S3C).

Collectively, these data suggest that developmental OTR expressions differ quantitatively in different cortical areas and even different layers within the same cortical region.

### Receptor down-regulation is the main mechanism of postnatal OTR reduction

Reduction of OTR expressing cells in the adult isocortex can be explained by either receptor down-regulation or programmed cell death during early postnatal development. For example, 40% of interneurons in the mouse cortex are eliminated during the postnatal period via programmed cell death ^28^. To understand the main mechanism dictating the transient nature of OTR expression, we crossed mice expressing Cre recombinase under the OTR promotor (OTR-Cre knock-in mice) with Cre dependent reporter mice (Ai14) that express the tdTomato fluorescent reporter (Figure 4A). The presence of Cre, even if transient as seen during developmental periods, leads to permanent expression of tdTomato (Figure 4A). If OTR (+) cells were undergoing cell death, we would expect to see a reduced number of tdTomato (+) cells in the adult brain. On the other hand, if OTR is simply down-regulated but the cells remain, tdTomato (+) cell density should not decrease in adulthood. When we quantified cortical tdTomato (+) cells from OTR-Cre:Ai14 mice using the dqBrain method for OTR-Venus mice, we observed that the average density of tdTomato (+) cells began to plateau at around P21 without any reduction in the adult stage at P56 (Figure 4B-C). Rather, OTR density continued to increase slightly between P21 and P56 largely because of the developmental accumulation of tdTomato within cells leading to slightly higher cell counting in later ages. In summary, this data suggests that developmental regulation of OTR in the isocortex is mainly driven by receptor down-regulation, not by cell death.

**Figure 4.**
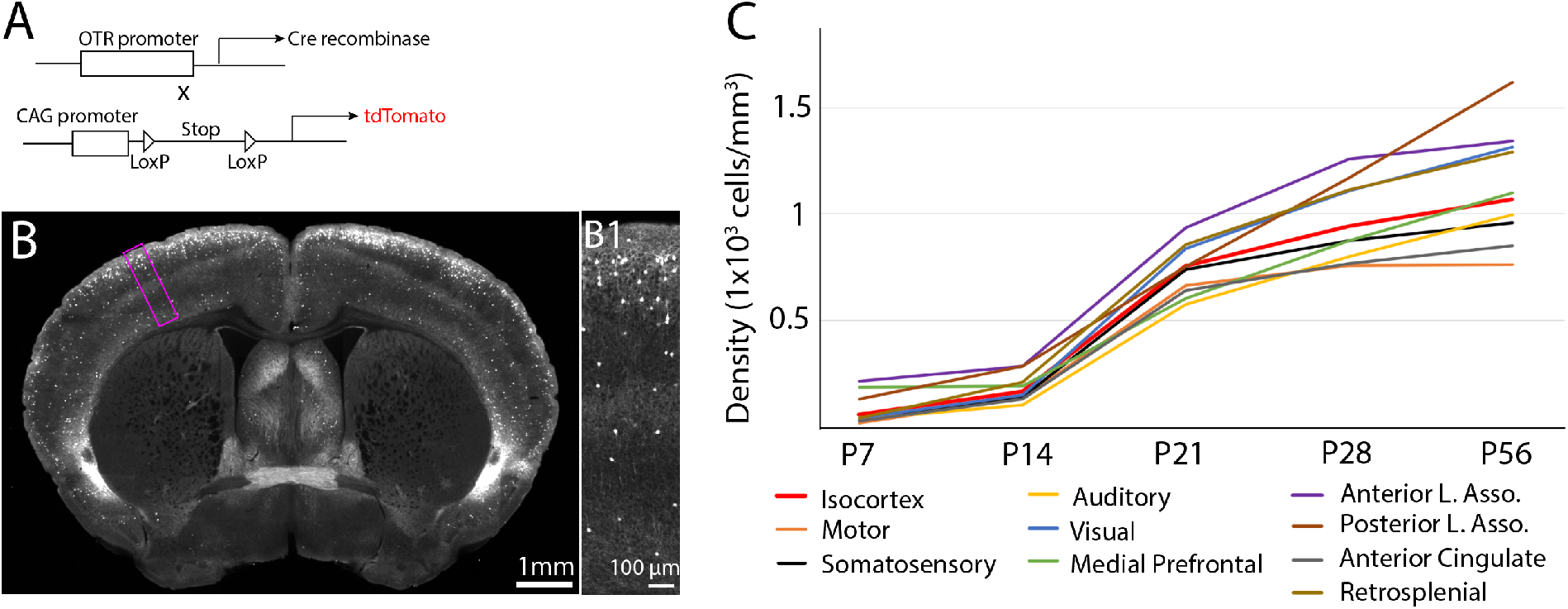
OTR down regulation is largely driven by receptor down-regulation. (A) Experimental design to permanently label transient OTR positive neurons by crossing OTR-Cre with Cre dependent reporter mice (OTR-Cre:Ai14). (B) Example of an adult OTR-Cre:Ai14 brain. (B1) High magnification image of purple boxed area in (B). Note the abundant tdTomato positive cells in the upper layer from the developmental labeling. (C) Average density of tdTomato (+) cells in different isocortical regions at different developmental time points.

### Layer specific cell type composition of OTR neurons in the isocortex

Neurons in the mouse isocortex are composed of non-overlapping glutamatergic (excitatory) and GABAergic (inhibitory) neurons with roughly a 4:1 ratio ^29^. OTR is known to be expressed in both glutamatergic and GABAergic cortical neurons ^14,27,30^. In order to determine the cell type of OTR expressing cortical neurons during postnatal development, we performed immunohistochemistry against GAD67, a marker for GABAergic neurons, in OTR-Venus mice at P21 and P56 (N = 3 mice per age, 3 representative sections per brain region; Figure 5). We examined the medial prefrontal cortex, the somatosensory cortex, and the visual cortex at three different cortical layers; upper layer for layer 1 – 3, deeper layer for layer 4 – 6a, and layer 6b (Table1). We found that the minority of OTR-Venus cells (at around 20%) in both upper and deeper layers are GABAergic in both ages (Figure 5, Table 1). In contrast, the majority of OTR-Venus cells in the deepest cortical layer (layer 6b) were GABAergic in both ages (Figure 5, Table 1). Interestingly, a previous study showed that these deep layer OTR positive neurons are mostly long-range projecting somatostatin neurons ^31^. There was no noticeable difference of OTR neuronal subtype composition in the isocortex between P21 and P56 (Table 1).

**Figure 5.**
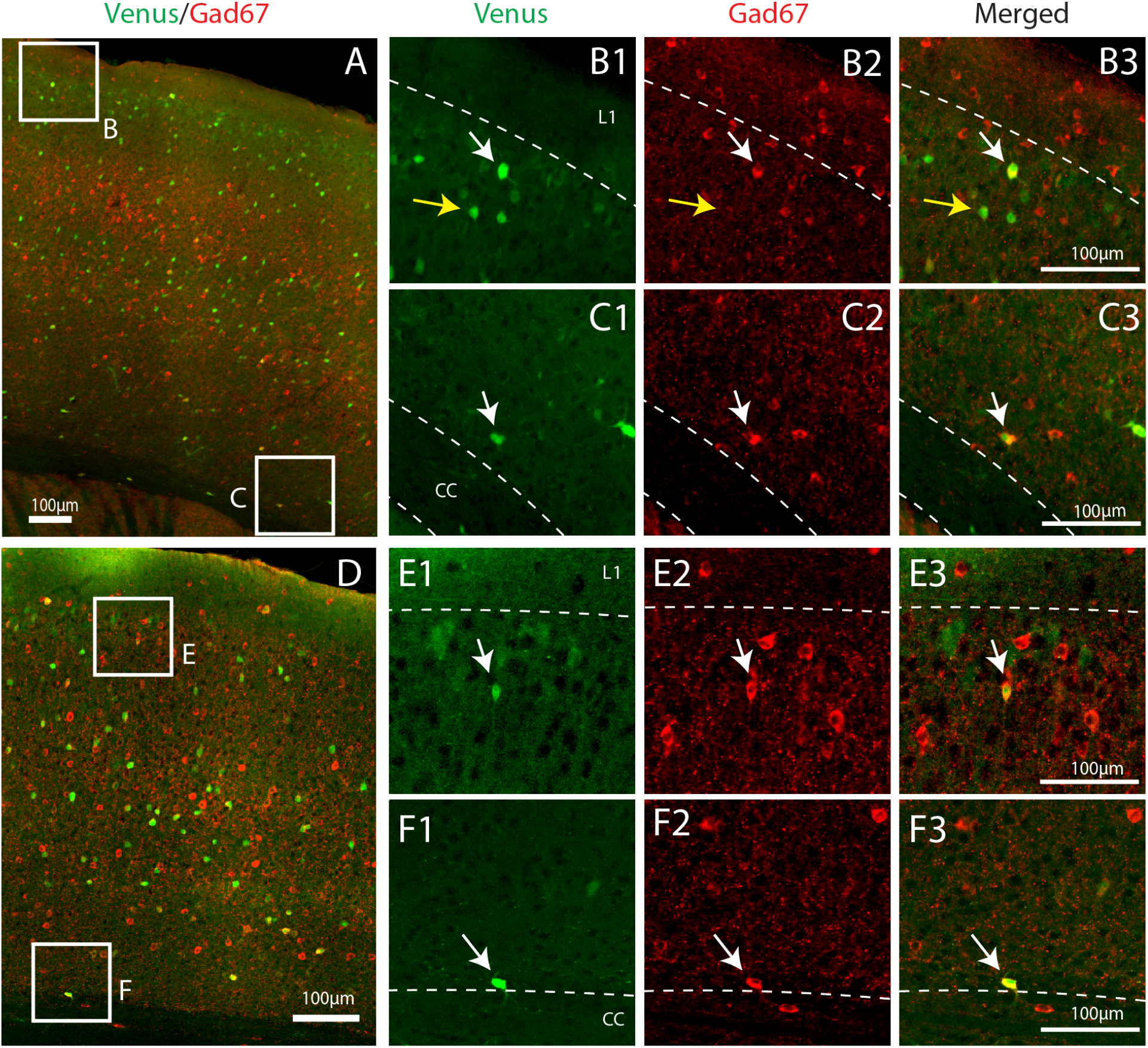
OTR cell types in the cortex. (A-E) Gad67 antibody immunohistochemistry staining (red) from motor-somatosensory cortical area at around bregma A/P = −0.7 mm from P21 (A-C) and P56 (D-E) OTR-Venus mice (green). (B-C, E-F) from the upper layer (B, E) and the layer 6b (C, F). white arrows for Venus (+) cells co-expressing Gad67, and yellow arrows for Venus (+) cells without Gad67 colocalization. L1 in (B, E) = layer 1, cc in (C, F) = corpus callosum.

**Table 1.** Gad67 colocalization with cortical OTR-Venus neurons. Data from the medial prefrontal cortex (at around Bregma A/P:+1.6), the somatosensory cortex (at around Bregma A/P:−1.0), and the visual cortex (at around Bregma A/P:−3.5). Data presented as percentage of colocalized cells (OTR and Gad positive cells/total OTR positive cells) in each brain region.

|  | **P21** |  |  | **P56** |  |  |
| --- | --- | --- | --- | --- | --- | --- |
| **Brain area** | **Upper layer** | **Deeper layer** | **Layer 6b** | **Upper layer** | **Deeper layer** | **Layer 6b** |
| **Medial prefrontal cortex** | 17%<br>(99/576) | 20%<br>(123/619) | 56%<br>(32/56) | 24%<br>(78/326) | 29%<br>(123/421) | 62%<br>(16/26) |
| **Somatosensory cortex** | 13%<br>(156/1191) | 19%<br>(301/1581) | 73%<br>(36/49) | 13%<br>(120/927) | 17%<br>(241/1399) | 74%<br>(38/51) |
| **Visual Cortex** | 18%<br>(123/671) | 15%<br>(214/1404) | 92%<br>(43/46) | 19%<br>(131/691) | 19%<br>(342/1809) | 86%<br>(40/46) |

### Developmental expression of OTR neurons in subcortical brain regions

Kinetics of neural circuit maturation vary significantly between different brain regions ^32^. Since OTR is widely expressed in different brain regions outside of the cortex, we sought to find whether these brain regions undergo similar expression trajectory to the isocortex in postnatally developing brains. We first examined ten large brain regions (the olfactory area, the hippocampal area, the striatum, the pallidum, the cerebellum, the thalamus, the hypothalamus, the midbrain, the pons, and the medulla) based on the Allen Brain Atlas ontology ^22^. The olfactory areas express OTR at the highest levels (Purple line in Figure 6A, Movie S1-5) as exemplified by a very high OTR density in the anterior olfactory nucleus (Figure 6B). In contrast, the cerebellum and the thalamus showed the lowest OTR densities (grey and yellow lines in Figure 6A, respectively) with a few noticeable exceptions including relatively high expression in the paraventricular thalamus (Figure 6E). There are also several noticeable subcortical areas with strong expression including the magnocelluar nucleus (also called “magnocellular preoptic area”), a part of the basal forebrain area (Figure 6C). We also observed prominent expression in specific hippocampal areas including the subiculum (Figure 6F). All areas except the hypothalamus reached their peak OTR densities at P21 with slight decrease in adulthood (Figure 6A). Interestingly, we observed continued increase of OTR in many hypothalamic nuclei until adulthood including the ventral medial hypothalamus ventral lateral, which matched previous OTR binding assays in rats ^33^(Figure 6A, D). A detailed list of OTR cell density across all brain regions at different ages can be found in Table S1.

**Figure 6.**
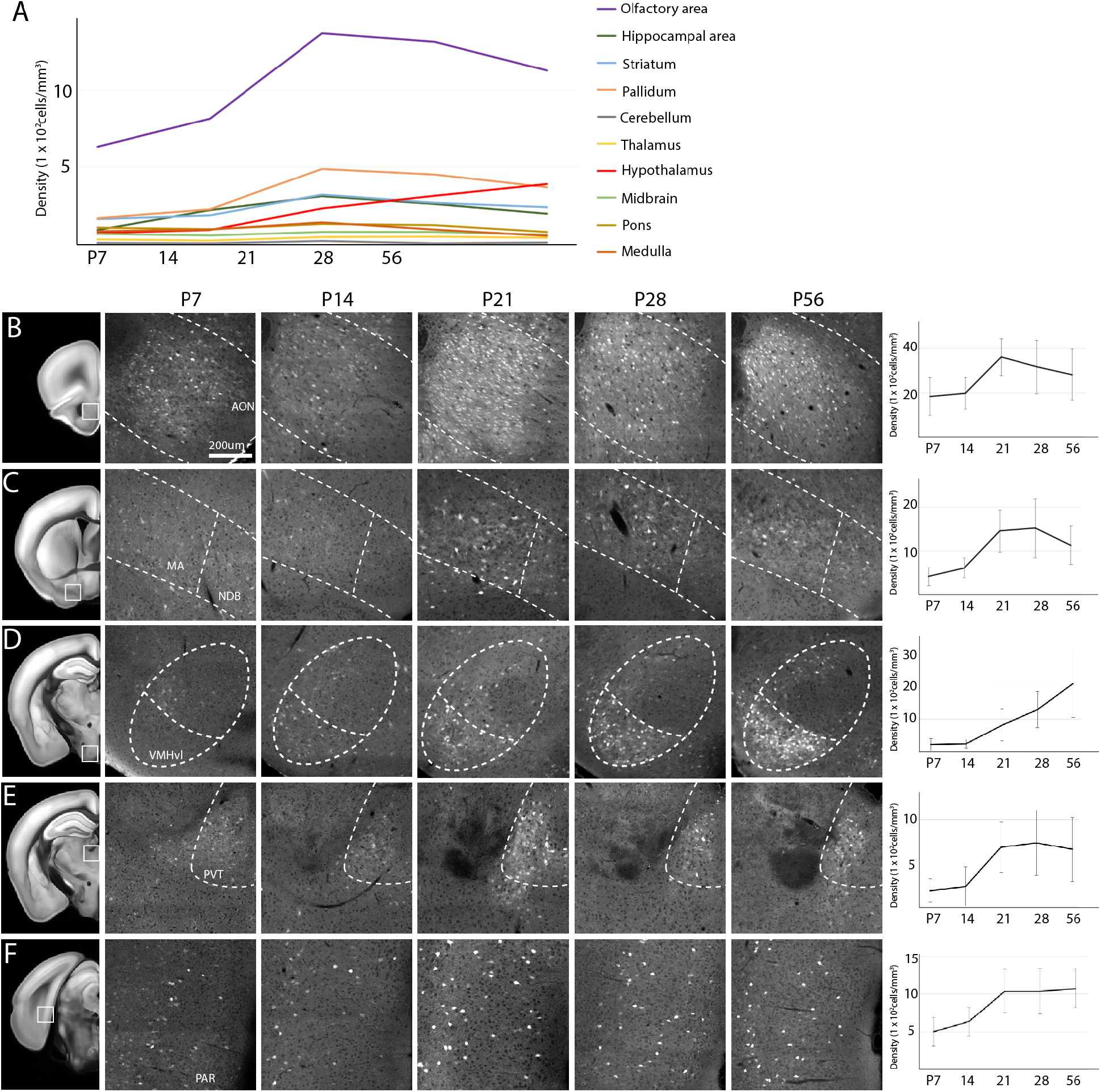
Temporal expression pattern of OTR neurons in other cortical and subcortical regions. (A) Average density of OTR neurons in ten different major sub-regions of the brain at postnatal development periods. (B-F) Notable brain regions with different temporal expression patterns. The first column highlights anatomical region of interest with white boxes in the adult reference brain. Mid columns represent zoomed-in picture of highlighted brain regions at different ages between P7 – P56. The last column is for the OTR (+) cell density measurement of the selected region (mean ± standard deviation). (B) The anterior olfactory nucleus (AON). (C) The magnocellular regions (MA) and the nucleus of diagonal band (NDB) in the basal forebrain area. (D) The ventral medial hypothalamus ventral medial (VMHvl) in hypothalamic areas. (E) The paraventricular thalamus (PVT) in thalamus. (F) The presubiculum (PRE) as a part of the retrohippocampal region.

### Sexual dimorphism of OTR expression

OTR is expressed in a sexually dimorphic manner as a part of neural circuit mechanism to generate behavioral differences in males and females ^34,35^. Therefore, we examined OTR-Venus mice (N=5 in each male and female brains at different ages) to determine if there were any regions showing strong sexual dimorphism. Across the entire brain region throughout the postnatal development, we found significant sexual dimorphism in two hypothalamic regions (Figure 7). The ventral premammillary nucleus showed significantly higher OTR expression in males compared to females between P14 and P56 (Figure 7A). In contrast, the anteroventral periventricular nucleus (AVPV) near the medial preoptic area showed higher OTR expression in females than males at P56, but not before (Figure 7B). A recent study identified abundant estrogen-dependent OTR expressing cells in the AVPV, co-expressing estrogen receptor in female mice ^36^. This result suggests a potential role of OTR in sexual behavior ^36–38^.

**Figure 7.**
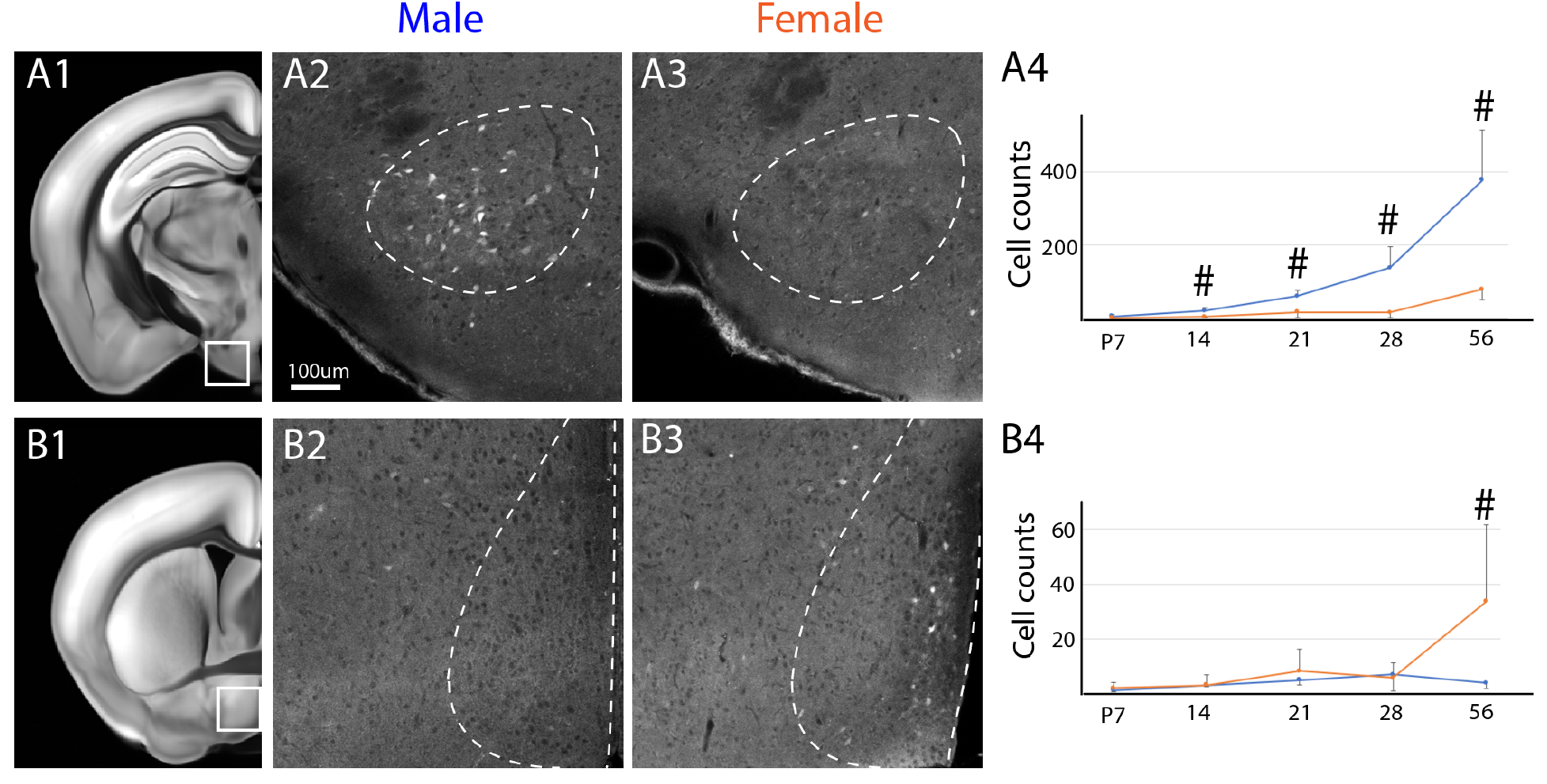
Sexually dimorphic expression of OTR neurons. (A) The ventral premammillary nucleus (PMv) showed significantly higher density of OTR cells in males than females from P14. (B) In contrast, the anterioventral periventricular nucleus (AVPV) near the medial preoptic nucleus showed significantly higher OTR cells in females than males at P56 but not before. The first column is to highlight anatomical regions of interest in an adult reference brain. 2^nd^ and 3^rd^ columns are zoomed-in pictures from P56 adult male and female OTR-Venus brains, respectively. The last column is density measurement over time (mean ± standard deviation). # denotes statistically significant data with false discovery rate less than 0.05 after multiple comparison correction.

### Web based resource sharing

Our high-resolution whole brain OTR expression dataset can serve as a resource for future studies examining how OTR regulates different neural circuits in postnatal development and adulthood. Moreover, our newly generated postnatal template brains can be used to map signals from different studies in the same spatial framework. To facilitate this effort, we created a website (http://kimlab.io/brain-map/OTR) to share our imaging data and other data resources from the current study. Data included in this paper can be easily visualized and downloaded from different web browsers including mobile devices. We highly encourage readers to explore this whole brain dataset on our website to investigate OTR expression in their regions of interest.

## Discussion

Here, we provide highly quantitative brain-wide maps of OTR expression in mice during the early postnatal developmental period and adulthood. We established new mouse brain templates at different postnatal ages and applied our dqBrain method to image and quantify fluorescently labeled signals at cellular resolution in postnatally developing brains ^23^. We found spatially and temporally heterogeneous developmental OTR expression patterns in different brain regions. Moreover, we found sexually dimorphic OTR expression in two hypothalamic regions. Lastly, our high-resolution imaging data is freely accessible via an online viewer as a resource for the neuroscience community.

OT signaling via OTR plays a pivotal role in postnatal brain development and is a key component of multi-sensory processing required to generate mature social behavior ^39,40^. Moreover, quantitative changes of OTR have been correlated with social behavioral variation in both normal and pathological conditions ^40,41^. For example, OTR expression levels within a brain region change based on early social experience ^42 43^. These findings suggest that OTR expression may be uniquely linked to the early postnatal development of social behavior. Thus, our data provides a quantitative understanding of OTRs developmental patterns in different neural circuits during critical periods of social behavior development.

We chose to use OTR-Venus mice to examine the whole brain OTR expression patterns throughout postnatal developmental periods after confirming that this reporter line provides a faithful representation of endogenous OTR expression using fluorescent *in situ* hybridization. The OTR-Venus expression patterns described here largely agree with previous histological studies focused on selected brain regions and/or ages ^6,7,21,33,40,44–47^. With its rapid protein maturation and decay half-life ^48,49^, Venus served as an ideal reporter protein for developmentally transient OTR expression in the entire brain which enabled us to circumvent laborious histological staining.

Our data driven approach uncovered quantitative insights about postnatal OTR expression. First, there are significantly heterogeneous spatial and temporal patterns of OTR expression across different cortical domains. For example, OTR expression emerged in visual-auditory cortices as early as P7 and then propagated to the somatosensory cortex at P14, reaching overall peak expression at P21. Since mice do not open their ears and eyes until about two weeks after birth ^50^, OTR expression in the visual-auditory areas precedes corresponding sensory inputs. Previous studies showed that OT signaling via OTR promotes synaptogenesis and facilitates synaptic maturation in postnatally developing brains ^10,12,51^. This evidence raises the possibility that early OTR expression may prime cortical areas for incoming sensory signals. Rapid increase of the OTR expression in P14 – 21 corresponds with the peak time of synaptic formation and maturation in rodent brains ^52,53^. Synaptic maturation patterns differ in cortical layers during early postnatal periods ^54,55^. For example, tactile stimulus specific activity pattern emerges in the superficial layers of the barrel cortex which is subsequently followed by deep layer maturation in mice ^55^. Interestingly, we found that OTR expressed more abundantly in the superficial layers at early postnatal time points (P7 and P14) followed by equal or relatively denser expression in the deep layer. This layer specific temporal cortical expression is mainly driven by transient OTR expression in the superficial layer at the early postnatal period between P14 – 21. Since this early postnatal period represents strengthening synaptic connections in the superficial layer 2/3 ^56^, transient OTR is ideally positioned to modulate synaptic maturation in the superficial layer. Second, we found that most subcortical regions also show their peak OTR expression at P21 followed by reduction into adulthood. This pattern agrees with previous observations that adult OTR patterns are established around three weeks postnatal age in mice ^7^. In contrast, the hypothalamic area showed a continuous increase into adulthood with sexually dimorphic expression of OTR in the PMv and AVPV, parts of the hypothalamic behavioral control column that generates sexually motivated behavior ^38^. This suggests that OTR in hypothalamic nuclei plays a role in generating sex-specific behavior during sexual maturation ^36,40,57^.

From a technical point of view, our dqBrain method is a significant departure from previous semi-quantitative histological methods. Our method provides a quantitative way to compare and contrast any fluorescently labeled signals in postnatally developing brains with various experimental conditions. Moreover, our newly established postnatal templates can help to map signals from other 3D imaging modalities (e.g., light sheet fluorescent microscopy) to age-matched spatial framework for quantitative comparisons. Previously, there has been significant effort to create common atlas framework to integrate findings from different studies in the adult mouse brain ^58–60^. Our postnatal template brains can serve as a common platform to study various signals from developing brains.

Lastly, our quantitative expression data with easy web based visualization provides a resource to examine OTR expression of any target brain region at different postnatal ages. Such open data sharing has proven to be useful in disseminating hard-earned anatomical data to the larger scientific community ^61,62^. In summary, we envision that our data will guide future circuit based investigation to understand the mechanism of oxytocin signaling in relationship with different behavioral studies in postnatally developing and adult brains.

## Material and Methods

### Animal

Animal procedures are approved by Florida State University, Tohoku University, and the Penn State University Institutional Animal Care and Use Committee (IACUC). Mice were housed under constant temperature and light condition (12 hrs light and 12 hrs dark cycle) and received food and water ad libitum. OTR-eGFP mice ^24^ were originally obtained from Mutant Mouse Resource and Research Center (MMRRC) with a mixed FVB/N x Swiss-Webster background strain. OTR-Venus mice ^20^ were originally produced and had their brains collected in Tohoku University (Nishmori Lab). Later, OTR-Venus line was imported to the Penn State University (Kim Lab). OTR-Venus brains used in the current study came from both Tohoku University and Penn State University. OTR-Cre line was originally established by Hidema et al., ^63^ and imported to the Penn State University via mouse rederivation. Both OTR-Venus and OTR-Cre mice are 129 × C57BL/6J mixed genetic background. OTR-Cre mice were then crossed with Ai14 (Jax: 007914, C57Bl/6J background) to generate OTR-Cre:Ai14 mice. All mouse lines were generated using continuously housed breeder pairs and P21 as the standard weaning date.

### Brain sample preparation, STPT imaging and related data analysis

Mice at various postnatal days were perfused by transcardiac perfusion with 0.9% NaCl saline followed by 4% paraformaldehyde in 0.1M phosphate buffer (PB, pH 7.4). Brains were further fixed overnight at 4°C and switched to 0.1M or 0.05M PB next day until imaging. Detailed procedure of STPT imaging was described previously ^23,26^. Briefly, fixed brains were embedded in oxidized agarose and cross-linked by 0.05M sodium borohydrate buffer at 4°C overnight to improve vibratome cutting during STPT imaging. For the STPT imaging, we used 910nm and 970nm to image OTR-eGFP and OTR^Venus/+^ mice, respectively. We acquired images at 1 μm (*x and y*) resolution in every 50 μm *z* section throughout the entire brain. For image registration to reference template brains, we used Elastix to register brains to age-matched reference template brains using 3D affine transformation with 4 resolution level, followed by a 3D B-spline transformation with 6 resolution level ^23^. We used a machine learning algorithm to detect fluorescently marked cells in serially collected 2D images. To convert the 2D counting to 3D counting, 2D cell counting numbers were multiplied by a 3D conversion factor (1.4) to estimate the total numbers of cells in each anatomical volume based on our previous calculation with cytoplasmic signals ^23^. To calculate the volume of each brain region, we registered age-matched template brains to each brain sample using Elastix. Then, voxel numbers of each anatomical label were multiplied by 20 × 20 × 50 μm^3^ (3D volume of anatomical voxel unit) to calculate volumes of each anatomical region ^23^. The 3D estimates of cell numbers in each anatomical region were divided by corresponding regional volume to generate density measurement per mm^3^ in each anatomical area (Table S1). All custom built codes were included in the previous publication ^23^.

### Statistical analysis

Density of fluorescently labeled cells in different anatomical regions including flatmap were presented as mean (Figure 3B-C, Figure 4C, Figure 6A, and Figure S3C) or mean ± standard deviation (Figure 6B-F and Figure 7A4 - B4). To identify sexual dimorphism, we performed statistical comparisons between males and females in OTR-Venus cell counts across different anatomical regions using open source statistical package “R”. We estimated our sample size using the power analysis as performed in our previous publication ^23^. When significance level (α < 0.05) and assumed effect size (0.85), we expected that over 80% of anatomical regions reach sufficient power with N = 5 samples per group. For statistical analysis between groups, we assumed the cell counts at a given anatomical area follow a negative binomial distribution and performed statistical analysis as described before ^23,26^. Once the p values were calculated, they were adjusted using false discovery rates with the Benjamini-Hochberg procedure to account for multiple comparisons across all ROI locations ^23,26^.

### Generating reference templates in different postnatal ages

All the work to generate the reference template brains at different ages was based on 20x downsized images in *x-y* dimension from the original scale, making each image stack at 20 × 20 × 50 x μm (*x,y,z*) voxel resolution. We picked the best-imaged brain with good right-left hemisphere symmetry (designated as a “template brain”) at each age and performed image registration using Elastix to register different age-matched brains to the template brain. Then, we averaged the transformed brains after the image registration to generate the averaged template brain at each age (Figure S2). We used either red or green channel images, or both from the same mouse acquired from the STPT imaging. To establish anatomical labels in averaged template brains, we used the image registration method to transform the adult atlas with anatomical labels to fit template brains at different ages. We used the common coordinate framework (CCF) brain and labels from Allen Institute for Brain Science as our initial atlas platform. Direct registration from the adult brain to averaged brains at each postnatal age worked well until the P14 brain due to similarities in postnatal brain morphologies, but not for P7 brains due to more embryonic brain-like shape. To circumvent this issue, we registered the adult CCF to the P14 template brain first, then the transformed CCF fit to the P14 brain was registered again to P7 (Figure S2). This sequential registration worked because the morphological differences between P7 and P14 were fewer than compared to those between P7 and the adult brain.

### Cortical Flatmap

We previously generated evenly spaced cortical bins to generate a cortical flatmap in an adult reference brain and devised a method to map detected signal in the flatmap ^23^. Here, we further generated superficial and deep layer cortical flatmaps. First, we created a binary file with layer 1 – 3 for superficial and layer 5 – 6 for deep layer across the entire isocortex. Second, we used the binary filter to remove unwanted cortical areas from the existing isocortical flatmap in order to create layer specific cortical flatmap. To quantify signals on flatmaps, we registered all samples to the reference brain with cortical area bins using Elastix and quantified target signals in each cortical bin as described in the STPT related data analysis above. We also performed reverse image registration to warp the adult reference brain to postnatal template brains in order to calculate the area of cortical bins at different ages. Then, we calculated densities in each cortical bin based on number of cells and area measurement in each bin. Lastly, the density was plotted in the cortical flatmap using Excel (Microsoft) and Adobe illustrator as described before ^23^.

### Single-molecule mRNA fluorescence *in situ* hybridization

Mice were deeply anesthetized using intraperitoneal injection of anesthesia (100 mg/kg ketamine mixed with 10 mg/kg xylazine). Then, the animal was decapitated with scissors, and the brain was immediately dissected out and immersed in Optimal Cutting Temperature (OCT) media (Tissue-Tek). The immersed brain was quickly frozen using dry ice chilled 2-methylbutane. The frozen brain was stored at-80°C until used. A cryostat was used to collect coronal brain sections at 10μm thickness. Sections were stored at-80°C, and *in situ* hybridization was performed within two weeks of sectioning. We used RNAScope detection kits (ACDBio) to detect and to quantify target mRNA at single-molecule resolution. We followed the manufacturer’s protocols with the exception that protease III (ACDbio, cat. no. 322340) was applied to tissue for 20 minutes. Probe-mm-Venus-C1 (ACDbio, cat. no. 493891) and Probe-mm-OTR-C2 (ACDbio, cat. no. 402651-C2) were used to detect Venus and OTR, respectively. Amp4 Alt A was used to detect OTR alone in red channel, and Alt C was applied to detect OTR and Venus in far red and red channel, respectively.

### Immunohistochemistry

*Sample preparation:* OTR^Venus/+^ mice of both sexes were collected at P21 and P56. Mice were deeply anaesthetized with intraperitoneal injection of the ketamine/xylazine mixture. Then, mice were transcardially perfused with 0.9% NaCl saline followed by 4% PFA. Whole heads were removed and post-fixed in the same fixative at 4°C for 3 days. Then, the brain was dissected out and sunk down in 30% sucrose in 1x PBS (pH7.4) solution at 4°C for cryoprotection. Cryoprotected brains were then frozen on dry ice and stored at −20°C until sectioning. 30 μm thick coronal sections were obtained using a freezing microtome (Leica). Sections were stored in a cryoprotectant solution (30% sucrose and 30% glycerol in 0.1M phosphate buffer) at-20°C until immunostaining. *Immunostaining:* All washes were performed for 10 min at room temperature with gentle rotation unless otherwise specified. Free floating sections were washed in 1x PBS three times followed by 1 hour of blocking with 1% donkey serum diluted in 1x PBS at room temperature. Slices were then incubated with a monoclonal primary antibody (mouse anti-GAD67, Millipore, cat.no. MAB5406, diluted 1:2000) in blocking buffer overnight at 4°C with gentle rotation. Following primary antibody incubation, the slices were washed in 1x PBS three times and incubated with secondary antibody (Donkey anti-mouse conjugated with Alexa 568, ThermoFisher, cat.no. A10037, diluted 1:500) for 1hr at room temperature. Three washes were performed in 0.05M phosphate buffer prior to mounting slices with vectashield mounting media (vector laboratories, cat.no. H-1500-10).

### Microscopic imaging and quantification

For both RNA *in situ* and immunostaining, BZ-X700 fluorescence microscope (Keyence) was used to image large areas using 20x objective lens with 2D image tile stitching. The sectioning function provided a deconvolution mechanism to capture sharply focused images. Images with large field of view were exported as Tiff files using the BZ-X analyzer software (Keyence). Image evaluation and cell counting was performed manually using the cell counter plug-in in FIJI (ImageJ, NIH) ^64^ All cell counting was done blindly by two independent experts. For the Venus and OTR colocalization in Figure 1J, Venus with more than 4 puncta from the RNA in situ was considered a Venus (+) cell. Images were acquired from cortical, amygdala, and hippocampal regions. In both OTR-Venus and OTR-Gad67 colocalization studies, two experts agreed over 95% of colocalization assessment. The final reported number is the averaged value from two expert’s counting.

## Acknowledgement

This publication was made possible by a NIH R01 MH116176 and Tobacco Cure Funds from the Pennsylvania Department of Health to Y.K., Strategic Research Program for Brain Sciences from Japan Agency for Medical Research and Development (AMED; 18dm0107076h0003, 2016-2020) to K.N. and S.H., JSPS Grant-in-Aid for Scientific Research (15H02442, 2015–2018) to K.N., NIH R01 MH114994 to E.A.D.H., T32 DC000044 to M.T. We thank Rebecca Betty for assistance in editing the manuscript.

Its contents are solely the responsibility of the authors and do not necessarily represent the views of the funding agency.

## Author Contribution

Project conceptualization: Y.K.; Brain Sample preparation and data acquisition: K.N., Z.T.N, U.C., M.T., S.H., K.N., E.H.; Data analysis: K.N., Z.T.N, A.R.W., Y.K.; Web visualization: D.J.V.; Manuscript preparation: K.N., Y.K with help of other authors.

---

**Figure S1.**
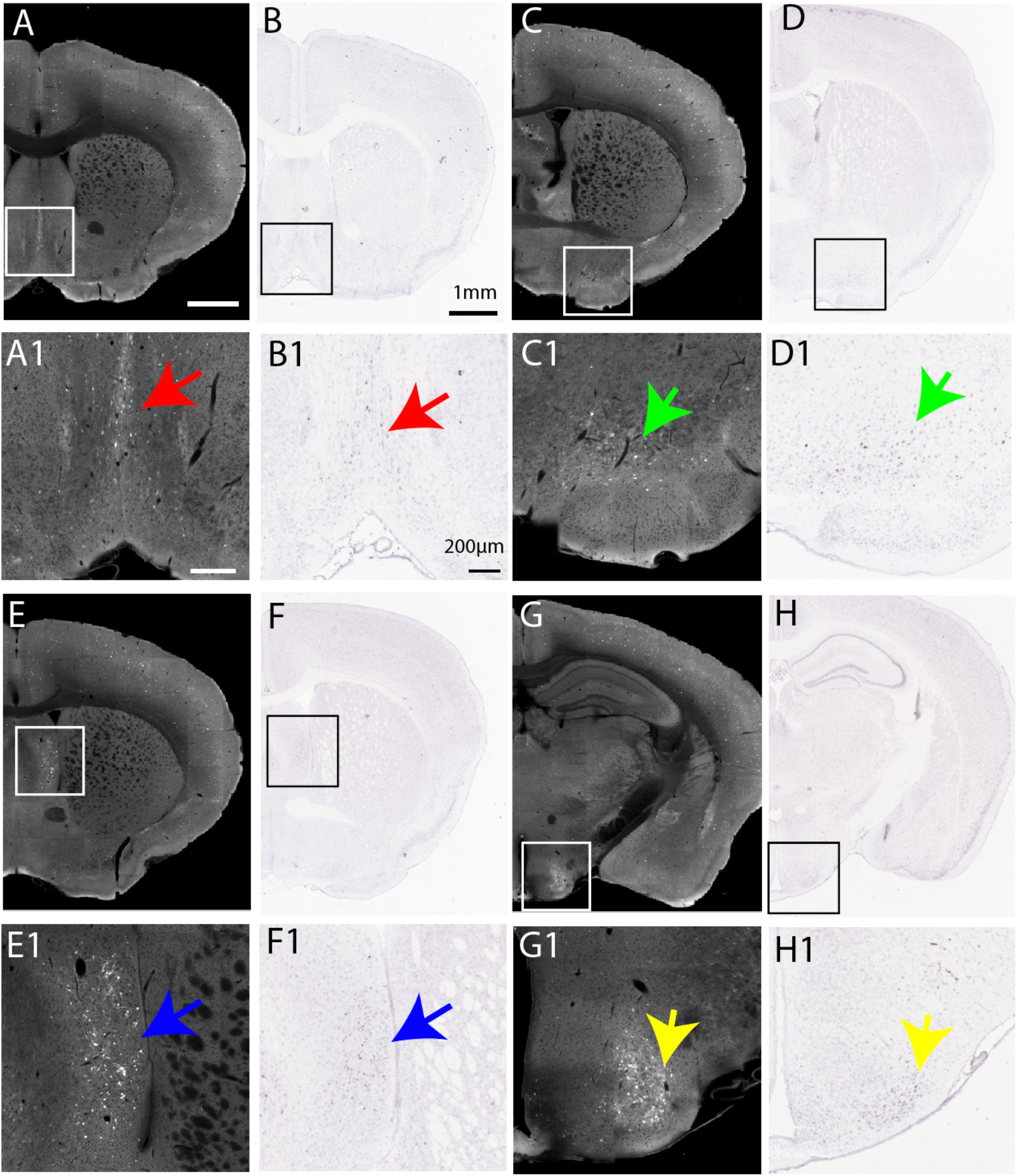
Comparison between Venus expression from OTR-Venus mice and OTR mRNA *in situ* at adult age. Venus expression from OTR-Venus (left column) and OTR *in situ* result (right column) from Allen in situ database (https://mouse.brain-map.org/experiment/show/75081001) in four different example areas: the medial septum (A-B), the nucleus of diagonal band (C-D), the lateral septum (E-F), and the ventral medial hypothalamus ventral lateral (G-H). The second row from each region is a zoomed-in view of the boxed areas in the first row of pictures. Note the matched pattern between OTR-Venus mice and endogenous OTR expression from the *in situ* data.

**Figure S2.**
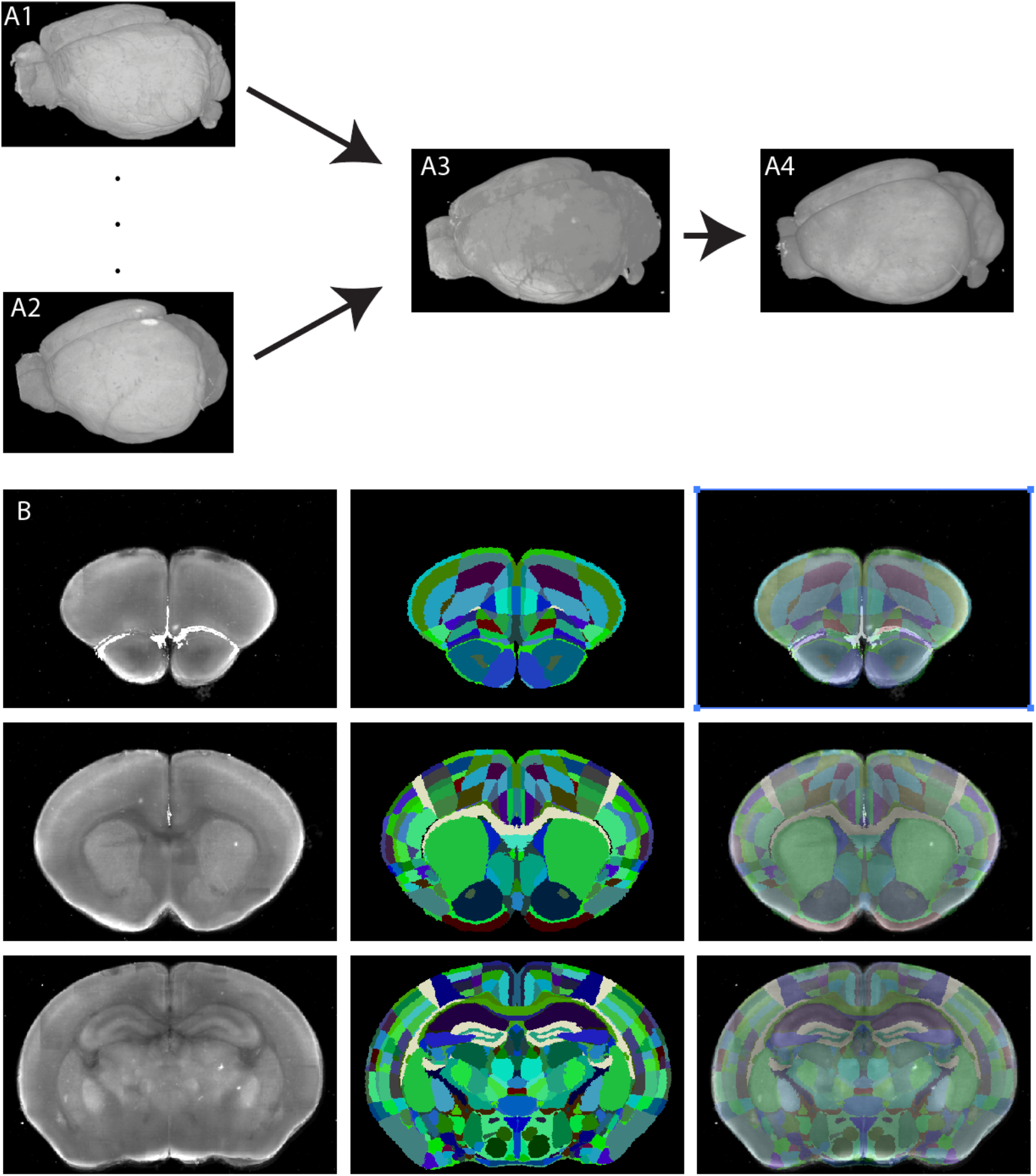
Generating age matched template brains and related anatomical labels. (A) Individual 3D brains (A1, A2) were registered to one best imaged sample (A3) from each age group. Registered brains were averaged to generate a template brain (A4) at each age. (B) Examples of anatomical labels from the P7 template brain. Row represents different areas in anterior and posterior axis. The first column is coronal view of the template brain, the second column for registered anatomical labels and the third row for the overlay between the template brain and labels.

**Figure S3.**
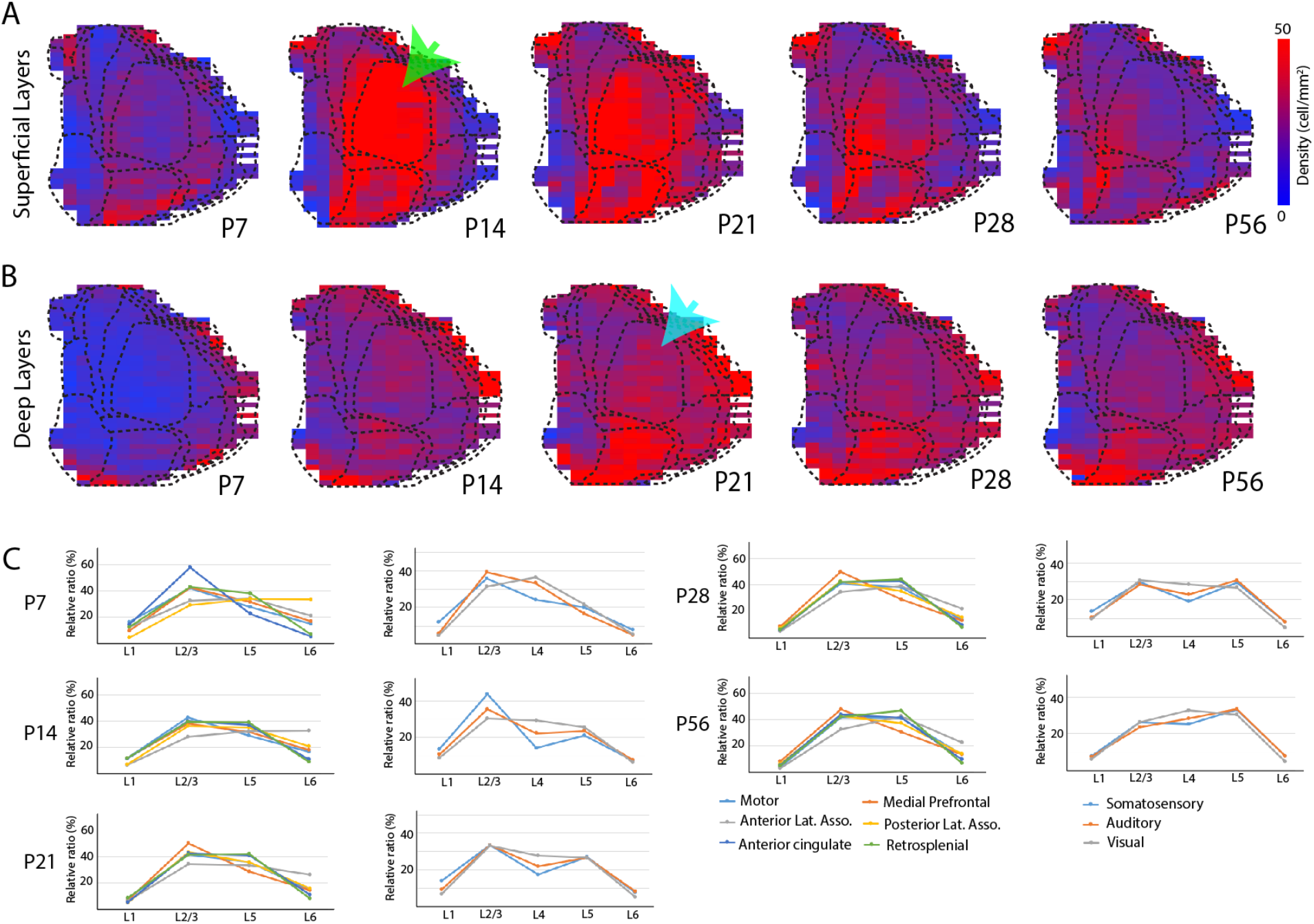
OTR expression in the layer specific cortical flatmap from OTR-Venus mice. (A-B) OTR-Venus expression patterns in the 2D cortical flatmap from superficial (A, layer 1-3) and deep (B, layer 5-6) layers at different postnatal ages. The heat map displays the visual representation of density. Note peak OTR density in the somatosensory cortex at P14 in the superficial layer flatmap (green arrow in A) while the peak at P21 in the deep layer flatmap (light blue arrow in B) for temporally heterogeneous OTR expression. (C) Relative densities of OTR-Venus cells across cortical layers in brain regions with motor and associative cortex (left column), and sensory cortex with layer 4 (right column). Density in each layer is normalized by total density of the whole layer in each brain region.

**Table S1. A list of OTR Densities across different brain regions at different postnatal ages.**

Column A: Acronym of brain regions, Column B: Full names, Column C-G: Average density (cell/mm^3^) at different postnatal ages. These columns are conditionally formatted with red color to highlight areas with high density. The heatmap color range between 0 (transparent) and 5000 (red). Column H-L: standard deviation at different postnatal ages.

**Movie S1-5: Quantitative OTR density mapping overlaid in age matched reference brains**.

Averaged OTR densities per evenly spaced and overlapping voxel (100 μm diameter sphere, 20 μm spacing between voxels) per postnatal age with the green heatmap to represent densities. The heatmap ranges between 0 (transparent) and 10 (green) cells/voxel. Left: overlay on the reference brain. Right: overlay on anatomical segmentations.

Movie S1 for P56, Movie S2 for P28, Movie S3 for P21, Movie S4 for P14, and Movie S5 for P7

